# Effects of Na^+^ channel isoforms and lipid membrane environment on temperature tolerance of cardiac Na^+^ current in zebrafish (*Danio rerio*) and rainbow trout (*Oncorhynchus mykiss*)

**DOI:** 10.1101/2020.11.20.391334

**Authors:** Jaakko Haverinen, Irina Dzhumaniiazova, Denis V. Abramochkin, Minna Hassinen, Matti Vornanen

## Abstract

Heat tolerance of heart rate in fish is suggested to be limited by impaired electrical excitation of the ventricle due to the antagonistic effects of high temperature on Na^+^ (I_Na_) and K^+^ (I_K1_) ion currents (I_Na_ is depressed at high temperatures while I_K1_ is resistant to them). To examine the role of Na^+^ channel proteins and the lipid matrix of the channels in heat tolerance of I_Na_, we compared temperature-dependencies of zebrafish (*Danio rerio*) and rainbow trout (*Oncorhynchus mykiss*) ventricular I_Na_, and I_Na_ generated by the cloned zebrafish and rainbow trout Na_V_1.4 and Na_V_1.5 Na^+^ channels in HEK cells. Whole-cell patch clamp recordings showed that zebrafish ventricular I_Na_ has better heat tolerance and slower inactivation kinetics than rainbow trout ventricular I_Na_. In contrast, heat tolerance and inactivation kinetics of zebrafish and rainbow trout Na_V_1.4 channels are similar when expressed in the identical plasma membrane lipid matrix of HEK cells. The same applies to Na_V_1.5 channels. Thermal adaptation of the ventricular I_Na_ is largely achieved by differential expression of Na^+^ channel alpha subunits: zebrafish which tolerate well high temperatures mainly express the slower Na_V_1.5 isoform, while rainbow trout which prefer cold waters mainly express the faster Na_V_1.4 isoform. Differences in elasticity (stiffness) of the lipid bilayer may be also involved in thermal adaptation of I_Na_. These findings suggest that both the protein component and its lipid bilayer matrix are involved in thermal adaptation of the voltage-gated Na^+^ channels and therefore in heart rate regulation under thermal stress in fish.

## INTRODUCTION

Temperature is a major environmental factor that has exerted a strong selective pressure on animal life forms. Temperature-driven evolution has led to significant variation in thermal tolerance of ectothermic vertebrates, including fishes (Beitinger, 2000; Johnston and Bennett, 2008). Some fish species are adapted to a relatively narrow range of temperatures such as the stenothermic teleosts of the Antarctic Ocean and the polar cod (*Boreogadus saida*) of the Arctic Ocean (Somero and DeVries, 1967; Beers and Sidell, 2011; Drost et al., 2016; Abramochkin et al., 2019). In temperate climates, fishes usually have a wider thermal tolerance range, e.g. the rainbow trout (*Oncorhynchus mykiss*) are cold-water fish which survive temperatures between 0 and 28°C (Hokanson et al., 1977; Beitinger, 2000). The zebrafish (*Danio rerio*), teleost fish of the lakes and rivers of South-eastern Asia, also tolerate a wide temperature range (7-41°C) but occupy warmer habitats than rainbow trout (Cortemeglia and Beitinger, 2005; López-Olmeda and Sánchez-Vázquez, 2011).

Despite considerable research efforts, the physiological basis for the high temperature tolerance of fishes and other ectotherms remains unresolved. Practically all major processes and functions of the animal body (e.g. circulation, sensory/motor functions, behaviour, metabolism, digestion, immune defence, reproduction) are affected by temperature and therefore more or less adapted to the habitat temperature of the animal (Angilletta Jr et al., 2002; Gracey et al., 2004; Podrabsky and Somero, 2004; Vornanen et al., 2005; MacMillan, 2019). Therefore, it is likely that when approaching the upper temperature tolerance limit of an animal several processes simultaneously weaken in the animal’s body, but the severity of their effect at the organismal level may vary and manifest with varying delays (Vornanen 2020). Alternatively, different processes/functions may have slightly different thermal optima or failure temperatures for example depending on the level of biological organization where the process/function appears (Lagerspetz, 1987; Clark et al., 2013). A fundamental question is: is there a single underlying process or interaction principle that is common to the thermal disorders of various body functions? Because similar thermal limitations appear to apply to unicellular organisms and Metazoa, it has been suggested the limiting processes of animal life are found at the molecular level (Tattersall et al., 2012).

Electrical excitability is a common process for most tissues of the animal body including nerves, skeletal muscles, heart, and smooth muscles. These tissues are responsible for almost all vital functions of the animal body like sensation, learning, behaviour, locomotion, blood circulation, digestion, and homeostasis of the body (Hille, 2001). Recently, we have gathered evidence showing that high temperatures can impair electrical excitability and therefore potentially limit the upper thermal tolerance of both ectothermic and endothermic animals (Vornanen, 2020). Electrical excitability or generation of propagating action potentials (APs) is the result of interaction between inward and outward directed flow of ions across the plasma membrane. In studies on thermal tolerance of electrical excitability we have used fish ventricular myocytes as a model system, since they are well suited for patch-clamp experiments. In a quiescent cardiac myocyte, there is a negative resting membrane potential (V_rest_), which is maintained by K^+^ efflux via the inward rectifier K^+^ current (I_K1_). For initiation and propagation of cardiac AP, V_rest_ must be depolarized to the threshold potential (V_th_) of an AP. This is accomplished by Na^+^ influx through the voltage-gated Na^+^ channels, which generate the sodium current (I_Na_). AP is initiated only when the charge transfer of the inward I_Na_ (the source current) exceeds the charge transfer of the outward I_K1_ (the sink current or resting membrane leak). At high temperatures, electrical excitability of fish ventricular myocytes may fail due to the mismatch between the source current (I_Na_) and the sink current (I_K1_) (Vornanen et al., 2014). Acute warming reduces charge transfer via the I_Na_, while K^+^ leak via the I_K1_ increases: I_Na_ fails to depolarize V_rest_ to the threshold potential of AP. At the level of a working heart this appears as atrioventricular block and depression of heart rate (Haverinen and Vornanen, 2020), which may eventually result in a collapse of cardiac output and thermal death of the fish.

The present study aimed to test the source-sink mismatch hypothesis with respect to the source current, I_Na_ (Vornanen, 2020). Given the differences in temperature tolerances between rainbow trout (a cold-active temperate species) and zebrafish (a warm-dwelling subtropical species) it was hypothesized that thermal tolerance of the zebrafish I_Na_ is higher than that of the rainbow trout I_Na_. This assumption was tested by comparing I_Na_ of zebrafish and rainbow trout ventricular myocytes at different temperatures. Another prediction of the source-sink hypothesis is that the channels that generate currents with slow gating kinetics can withstand high temperatures better than channels that produce currents with fast kinetics (Touska et al., 2018; Vornanen, 2020). This is based on the assumption that stiffer molecules have higher activation energy and slower kinetics, i.e. there is a trade-off between flexibility and thermal stability of the proteins (Somero, 1995; Zavodszky et al., 1998; Fields, 2001). I_Na_ with slow inactivation kinetics is able to provide more depolarising charge at high temperature than I_Na_ which is rapidly inactivated. To this end the inactivation kinetics of I_Na_ between rainbow trout and zebrafish ventricular myocytes were compared at different temperatures.

Finally, it was hypothesised that thermal stability and kinetics of fish I_Na_ depend in part on the biophysical properties of the lipid membrane into which they are embedded. To test this, the two main alpha subunits of zebrafish and rainbow trout Na^+^ channels were cloned and expressed in a mammalian cell line, the human embryonic kidney (HEK) cells. This allowed comparison of inactivation kinetics and thermal resistance of the Na^+^ channels in an identical membrane matrix.

## MATERIALS AND METHODS

### Animals

The wild-type zebrafish (*ab* strain, kindly donated by Dr. Maxim Lovat, Lomonosov Moscow State University) were raised and maintained at the animal facilities of Lomonosov Moscow State University according to the common practices (Westerfield, 2007). The rearing temperature of the fish was 28°C. Fish of either sex, about 1.5-year-old, were used for electrophysiological experiments (n=8). Zebrafish were killed by immersion in an ice-water bath and cutting of the spine. Rainbow trout (*Oncorhynchus mykiss*) were obtained from a local fish farm (Kontiolahti, Finland). Fish weighing 118.7 ± 20.6 g (n=11) of either sex, acclimated at 12°C for more than 3 weeks, were used in electrophysiological experiments and gene cloning. Trout were stunned by a quick blow to the head and killed by cutting of the spine immediately behind the head. The experiments conform to the “European Convention for the Protection of Vertebrate Animals used for Experimental and other Scientific Purposes” (Council of Europe No 123, Strasbourg 1985) and were authorized by the national animal experimental board in Finland (permission ESAVI/8877/2019).

### Cloning of SCN5LA and SCN4A Na^+^ channel genes of zebrafish and rainbow trout

Total cardiac RNA was extracted by TriReagent (Thermo Scientific, Vilnius, Lithuania) and the quality and quantity of RNA were determined by agarose gel electrophoresis and NanoDrop ND-1000 Spectrophotometer (NanoDrop Technologies, Wilmington, DE, USA), respectively. RNA (2 μg) was treated with RNase free DNase (Thermo Scientific) and converted to cDNA using SuperScript IV reverse transcriptase (Invitrogen) and oligo(dT) primers. 1 μl of the cDNA was used as a template in 25μl polymerase chain reaction (PCR) including a final concentration of 0.2 mM dNTP mix, 0.2 mM primers (synthesized by Invitrogen, Glasgow, UK) shown in Table 1, and 0.02 U/μl Phusion High Fidelity DNA polymerase (Thermo Scientific). Cycling conditions for all PCR-reactions were as follows: initial denaturation at 98 °C for 1 min, 35 cycles at 98 °C for 10 s, at 60 °C for 30 s and at 72 °C for 100-260 s (40 s/kb, for the length of the products see Table 1), followed by final extension at 72 °C for 5 min. The PCR products were separated on a 0.8% agarose gel, and the nucleotide chains were extracted from the gel using the GeneJET Gel Extraction Kit (Thermo Scientific). Overhang adenines were added to the 3′-ends of PCR products using Dynazyme II DNA polymerase (Thermo Scientific) and products were ligated into the pGEM-T Easy vector (Promega Madison, Wisconsin, USA). The inserts were digested from the pGEM-T Easy vector and directionally cloned into the expression vector. If the coding sequence of the SCN gene was amplified by PCR in two parts, the pieces were ligated into the expression vector to form the entire coding sequence. Coding sequences of the SCN5LA genes of rainbow trout (om for *Oncorhynchus mykiss*) and zebrafish (dr for *Danio rerio*) were cloned into the expression vector pcDNA3.1/Zeo(+) (Invitrogen). Several attempts to clone the full-length om/drSCN4A into the pcDNA3.1 failed. To overcome this problem, the high-copy number vector pcDNA3.1/Zeo(+) was converted to a low-copy number vector that was better tolerated by bacterial cells. To this end, the ColE1 origin was replaced by the pMB1 origin via digesting pcDNA3.1/Zeo(+) with BsmI and PvuI and replacing this fragment (bases 2768-4462) with the digested fragment from pBR322 (bases 1353-3733). The coding sequences of dr/omSCN4A were successfully cloned into this modified vector. Plasmids were isolated using PureLink HiPure Plasmid Midiprep Kit (Thermo Scientific) and sequenced by GATC Biotech AB (Köln, Germany). The fish SCN4A and SCN5LA genes are orthologous to the mammalian SCN4A and SCN5A genes. For simplicity, the protein names (Na_V_1.4 and Na_V_1.5) of the orthologous genes are used throughout the text.

**Table 1.**
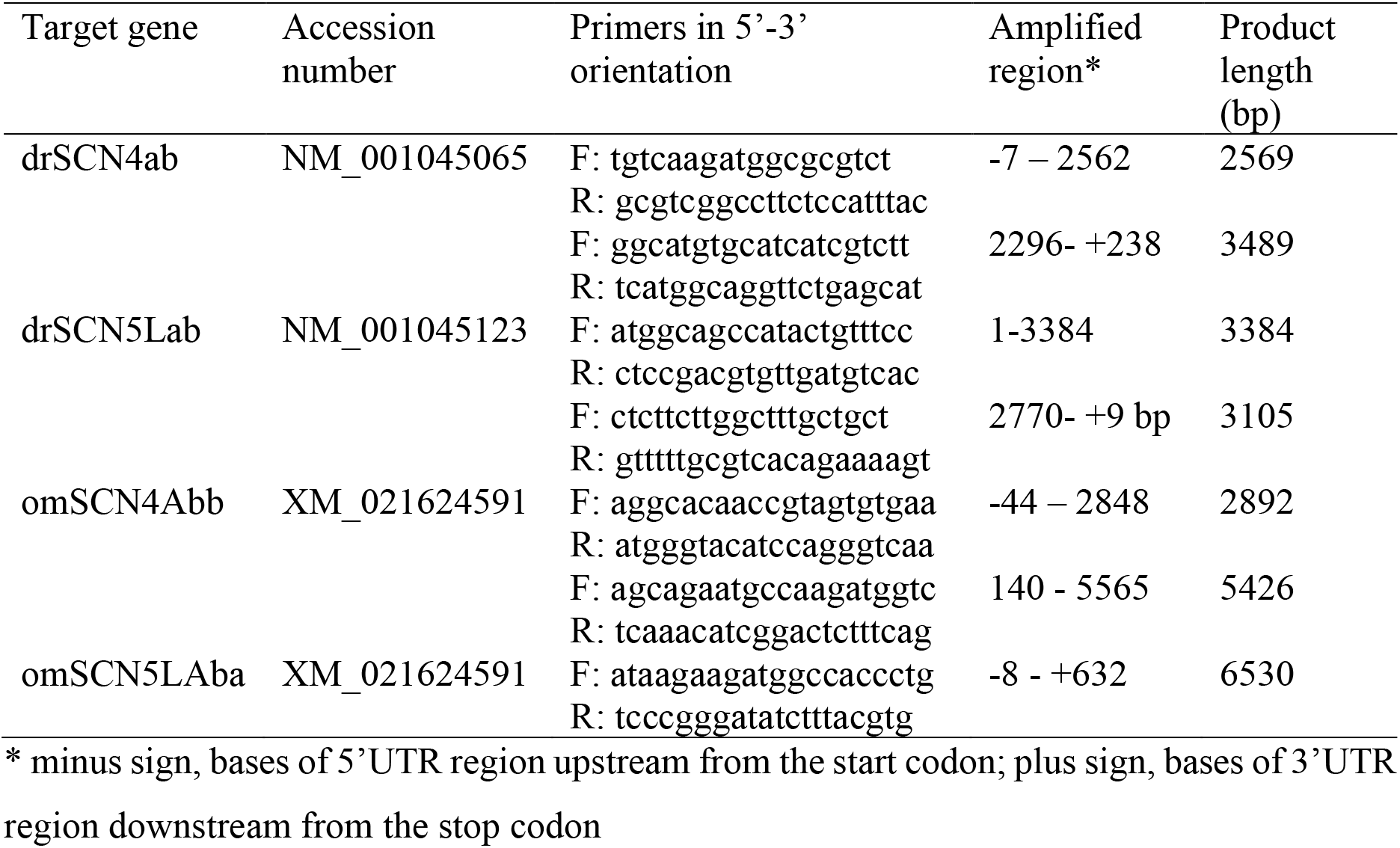
Primers used in cloning of the SCN genes of rainbow trout and zebrafish.

### Heterologous expression of SCN4A and SCN5LA genes

Human embryonic kidney (HEK293; ECACC) cells were grown at 28°C in DMEM (Biowest) supplemented with 10 % fetal bovine serum (FBS, sterile-filtered and heat inactivated; Sigma-Aldrich) and 100 U/ml penicillin and streptomycin (Sigma-Aldrich) at 37°C in 5% CO_2_ environment. HEK cells were transiently cotransfected with pEGFP-N1 (Clontech), and either drSCN4Ab, drSCN5LAb, omSCN4Abb or omSCN5LAba construct using TurboFect transfection reagent (Thermo Scientific). Cells were transfected for 16-18 hours at 37 °C after which fresh medium was changed to the plates and the cells were further incubated at 28 °C in 5% CO_2_ environment for at least 24 hours. The incubation at lower temperature increased the expression level of Na^+^ channels. Whole cell patch-clamp experiments were conducted 48–72 h after transfection.

### Patch clamp measurements of Na^+^ current in fish cardiac myocytes and HEK cells

The procedure of isolating zebrafish cardiomyocytes was essentially similar to the original method of isolating crucian carp (*Carassius carassius*) cardiac cells (Vornanen, 1997), but scaled down to the size of small zebrafish hearts and using slightly higher enzyme concentrations (1 mg/mL collagenase Type IA and 0.67 mg/mL Trypsin IV both from Sigma). A blunt-ended syringe needle (34 gauge, TE734025, Adhesive Dispensing Ltd, UK) cannula was inserted via *bulbus arteriosus* into the ventricle and secured in place with a fine thread. The heart was perfused first with Ca^2+^-free solution for 5 min and then with the same solution but with added hydrolytic enzymes together with fatty acid-free serum albumin (1 mg ml^−1^, Sigma) for 25-30 minutes. Myocytes were stored at 5°C and used on the same day as they were isolated. The Ca^2+^-free solution contained (in mmol l^−1^) 100 NaCl, 10 KCl, 1.2 KH_2_PO_4_, 4 MgSO_4_, 50 taurine, 20 glucose and 10 HEPES at pH 6.9 (adjusted with KOH at 20°C). From rainbow trout heart, myocytes were obtained using essentially the same procedure, but with lower enzyme concentrations (0.75 mg/mL collagenase Type IA and 0.5 mg/mL Trypsin IV) and shorter digestion time (10-12 minutes).

The whole-cell voltage clamp recording of I_Na_ was performed using an Axopatch 200B or an Axopatch 1-D amplifier (Molecular Devices, CA, USA) as previously described in detail (Haverinen et al., 2018). Cardiac myocytes or coverslips containing cultured HEK cells were placed in a small chamber with a continuous flow of K^+^-based external saline solution containing (in mmol l^−1^) 150 NaCl, 5.4 KCl, 1.8 CaCl_2_, 1.2 MgCl_2_, 10 glucose, 10 HEPES, with pH adjusted to 7.7 at 20°C with NaOH. The temperature of the saline solution was set using a Peltier device (CL-100, Warner Instruments, CT, USA or TC-10, Dagan, Minneapolis, USA) and monitored continuously with thermistors placed close to the cells. Patch pipettes with a resistance of 1.5-2.5 MΩ were drawn from borosilicate glass (Garner, Claremont, CA, USA) and filled with Cs^+^-based electrode solution containing (in mmol l^−1^): 5 NaCl, 130 CsCl, 1 MgCl_2_, 5 EGTA, 5 Mg_2_ATP and 5 HEPES with pH adjusted to 7.2 with CsOH. Current amplitudes were normalized to the capacitive cell size to obtain current density (pA·pF^−1^).

During I_Na_ recording, myocytes were superfused with a Cs-based low-Na^+^ external saline solution, which contained (in mmol l^−1^) 20 NaCl, 120 CsCl, 1 MgCl_2_, 0.5 CaCl_2_, 10 glucose and 10 HEPES at pH 7.7 (adjusted with CsOH at 20°C) (Haverinen and Vornanen, 2004). In experiments with cardiac myocytes, nifedipine (10 μmol l^−1^, Sigma) was included to block I_Ca_. The low concentration of Na^+^ outside the cell reduced the driving force for Na^+^ influx and made it possible to achieve stable and complete voltage control of the I_Na_. Approximately 80% of the series resistance was compensated when recording fast and large I_Na_. I_Na_ was leakage-corrected using the P/N procedure of the Clampex 9.2 software. For the determination of current-voltage (I-V) relationship, I_Na_ was elicited from the holding potential of −120 mV with 60 ms depolarizing pulses (range −100 - +70 mV) at the frequency of 1 Hz (Fig.1A). The voltage dependence of Na^+^ channel conductance was calculated from the I-V recordings using the equation (*G*_*Na*_ = *I*_*Na*_ /(*V* − *V*_*rev*_), where *G*_*Na*_ is the Na^+^ conductance of the membrane, I_Na_ is the peak current at a given membrane potential (*V*) and *V*_rev_ is the reversal potential of I_Na_. The steady-state (SS) voltage dependence of activation was obtained by plotting the normalized conductance (*G*/*G*_max_) as a function of membrane potential and fitting it into the equation of Boltzmann distribution 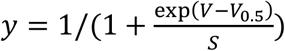 where *V* is membrane potential, *V*_0.5_ the midpoint potential and S the slope of the curve. SS inactivation was determined using a two-step protocol where a 300 ms conditioning pulse to potentials between −110 mV and −20 mV was followed by a 60 ms test pulse to −20 mV (Fig. 3a). The normalized test pulse currents (*I*/*I*_max_) were plotted as a function of membrane potential and fitted to the Boltzmann function with a negative slope (−S).

**Fig 1.**
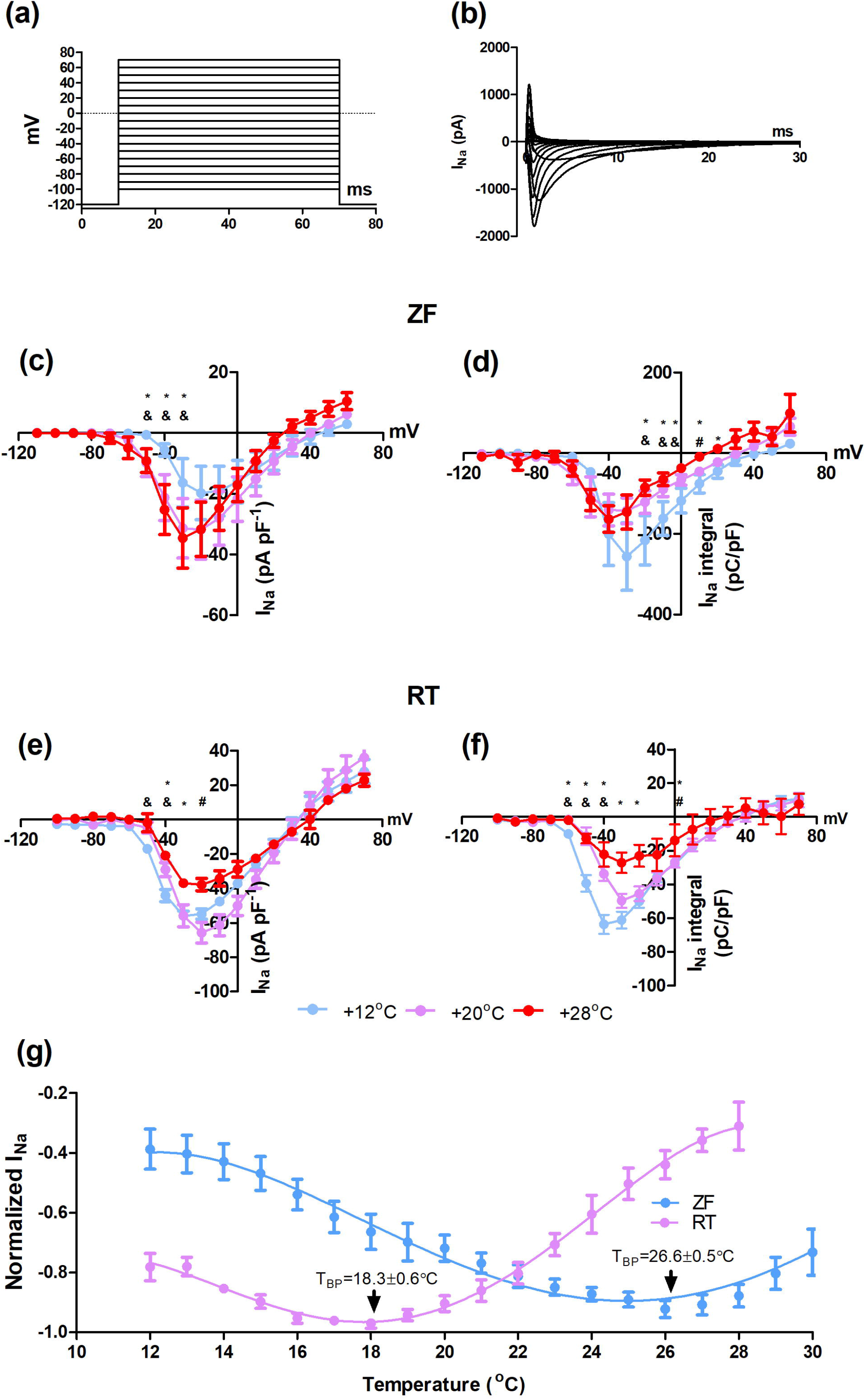
Voltage- and temperature-dependence of current density and charge transfer of I_Na_ in zebrafish and rainbow trout ventricular myocytes. (a) The voltage protocol used to elicit I_Na_. (b) A representative recording from a zebrafish ventricular myocyte indicating tracings of I_Na_ at different membrane voltages. (c-f) Mean results (± SEM) for current-voltage relationship and charge transfer of I_Na_ at three different temperatures for zebrafish (n=47) (c, d) and rainbow trout (n=33) (e, f) ventricular myocytes. (g, h) Current density of I_Na_ in heat ramp experiments with zebrafish (n=6) (g) and rainbow trout (n=5) (h) ventricular myocytes. Statistically significant differences (p<0.05) are shown as follows: * 12°C vs. 28°C, & 12°C vs. 20°C, # 20°C vs. 28°C.

The time constant of I_Na_ inactivation at different membrane potentials (−30 mV to 0 mV) was derived by fitting the decay phase of I_Na_ using the double exponential function of the Chebyshev transformation procedure of the Clampfit 10.3 software package. The amplitude of the fast component (τ_f_) was over 90% of the current. The slow component could only be reliably determined at the voltages at which I_Na_ was close to its peak amplitude. Therefore, only the results of τ_f_ are reported.

All mentioned properties of I_Na_ were analyzed in native trout or zebrafish myocytes and HEK cells expressing Na_V_1.4 or Na_V_1.5 channels at 3 different temperatures: 12°C, 20°C and 28°C. However, additional experiments to estimate the dynamics of the temperature dependence of I_Na_ were done using the “heat ramp” protocol. In these experiments I_Na_ was elicited by repetitive depolarizations to −20 mV and the temperature was steadily raised from 12°C to 28°C (up to 39°C in the case of native zebrafish myocytes).

### Statistics

Statistical analyses were performed using SPSS (IBM; version 25). One-way ANOVA (with Tukey’s or Dunnett’s T3 *post-hoc* test) and unpaired t-test were used to compare normally distributed data with homogenous variances. If the assumptions of parametric tests were not met, non-parametric Kruskal-Wallis and *post-hoc* Mann-Whitney *U* tests were used. A *p* < 0.05 was considered to show a statistically significant difference between means.

## RESULTS

### Temperature-dependence of I_Na_ density and charge transfer in zebrafish and rainbow trout ventricular myocytes

Temperature-dependency of ventricular I_Na_ was markedly different between zebrafish and rainbow trout myocytes as shown by the current-voltage (I-V) relationships (Fig 1). In zebrafish myocytes, an acute rise of temperature from 12°C to 20°C resulted in a significant increase in I_Na_ density at voltages between −50 and −30 mV (Fig. 1c). A similar difference was detected in I_Na_ density between 12°C and 28°C (p<0.05). However, the current densities at 20°C and 28°C were almost identical suggesting plateauing of I_Na_ in this temperature range (Fig 1c). In rainbow trout myocytes, warming from 12°C to 20°C increased I_Na_ density at voltages from −30 mV to −10 mV, but further warming to 28°C caused a strong depression of I_Na_ (Fig. 1e). Although the density of the zebrafish I_Na_ was higher at 20°C and 28°C than at 12°C the charge transfer (integral of I_Na_) was significantly smaller than at 12°C (Fig. 1d). In trout ventricular myocytes the integral of I_Na_ decreased with increasing experimental temperature (Fig. 1f). Notably, charge transfer/I_Na_ density ratio, i.e. depolarizing power, was 4.5-22.2 times higher (at −20 mV) for zebrafish than rainbow trout I_Na_ at all experimental temperatures (0.6, 0.7, and 0.9 V/A for rainbow trout and 2.7, 4.5 and 20.0 V/A for zebrafish at 28°C, 20°C and 12°C, respectively).

Temperature tolerance of peak density and charge transfer of I_Na_ (at −20 mV) were further studied using acute heat ramps. To this end, the cells were warmed from 12°C to their upper thermal tolerance limit while I_Na_ was elicited by depolarization from −100 mV to −20 mV for every second (Fig. 1 g). Consistent with the I-V data, there was a dramatic interspecies difference in temperature-dependence of the peak I_Na_ (Fig. 1g). Initially, when the temperature was raised above 12°C, the density of I_Na_ increased in both species. However, the breakpoint temperature (*T*_BP_, the temperature above which the peak I_Na_ started to decrease steadily) was much lower for rainbow trout I_Na_ (18.3±0.6°C, n=13) than for zebrafish I_Na_ (26.6 ± 0.5°C, n=16) (p<0.05).

Taken together, these findings indicate that the ventricular I_Na_ of rainbow trout heart is much less tolerant of high temperatures than the zebrafish ventricular I_Na_.

### Inactivation kinetics of I_Na_ in zebrafish and rainbow trout ventricular myocytes

Temperature-dependence of I_Na_ inactivation kinetics was measured in the voltage range from −30 to 0 mV at 12°C, 20°C and 28°C (Fig. 2). Kinetics of zebrafish or rainbow trout I_Na_ accelerated with increasing temperature and membrane depolarization. There was a striking difference in the rate of I_Na_ inactivation between trout and zebrafish ventricular myocytes (Fig. 2 a-f). At 12°C, 20°C and 28°C the time constant (τ) of I_Na_ inactivation (at 0 mV) was about 10.0, 6.3 and 4.0 times faster in rainbow trout ventricular myocytes in comparison to zebrafish ventricular myocytes (p<0.05).

**Fig. 2.**
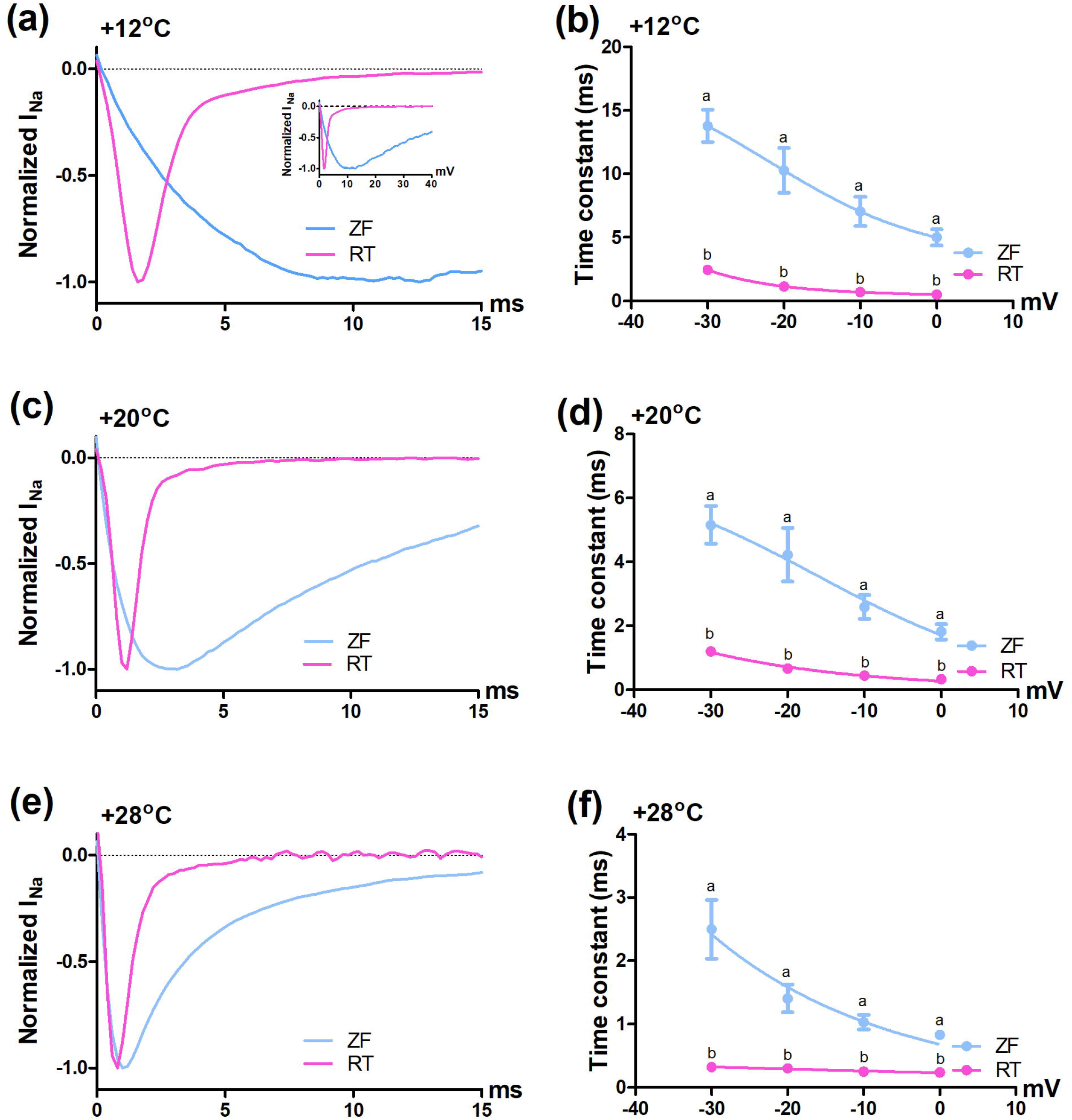
Temperature-dependence of inactivation kinetics of I_Na_ in zebrafish and rainbow trout ventricular myocytes. I_Na_ was elicited from a holding potential of −120 mV to −30 – 0 mV for 30 ms. (a, c and e) Representative tracings of I_Na_ at −20 mV at temperatures of 12°C (a), 20°C (c) and 28°C (e). (b, d and f) Means (± SEM) of inactivation time constant (τ) at 12°C (b), 20°C (d) and 28°C (f). The results are from 31 and 30 myocytes from zebrafish and rainbow trout, respectively. Error bars of trout data are smaller than the size of the symbols. Statistically significant differences (p<0.05) are shown as follows: * 12°C vs. 28°C, & 12°C vs. 20°C, # 20°C vs. 28°C.

### Steady-state activation and inactivation of I_Na_ in zebrafish and rainbow trout ventricular myocytes

Voltage-dependence of steady-state (SS) activation and inactivation of I_Na_ were studied at 12°C, 20°C and 28°C (Fig. 3). Voltage-dependence of I_Na_ SS inactivation was only slightly affected by acute temperature increases in zebrafish ventricular myocytes (Fig. 3c). Warming from 12°C to 20°C and further to 28°C slightly decreased the slope factor (p<0.05) (Table 2). Voltage-dependence of SS activation of I_Na_ was slightly more temperature-sensitive, since warming to either 20°C or 28°C shifted V0.5 to the left by almost 10 mV and decreased the slope factor (p<0.05) (Table 2, Fig. 3c). In trout ventricular myocytes warming from 12°C to 20°C and further to 28°C induced a slight positive shift of SS activation V_0.5_ (Table 3, Fig. 3d).

**Fig. 3.**
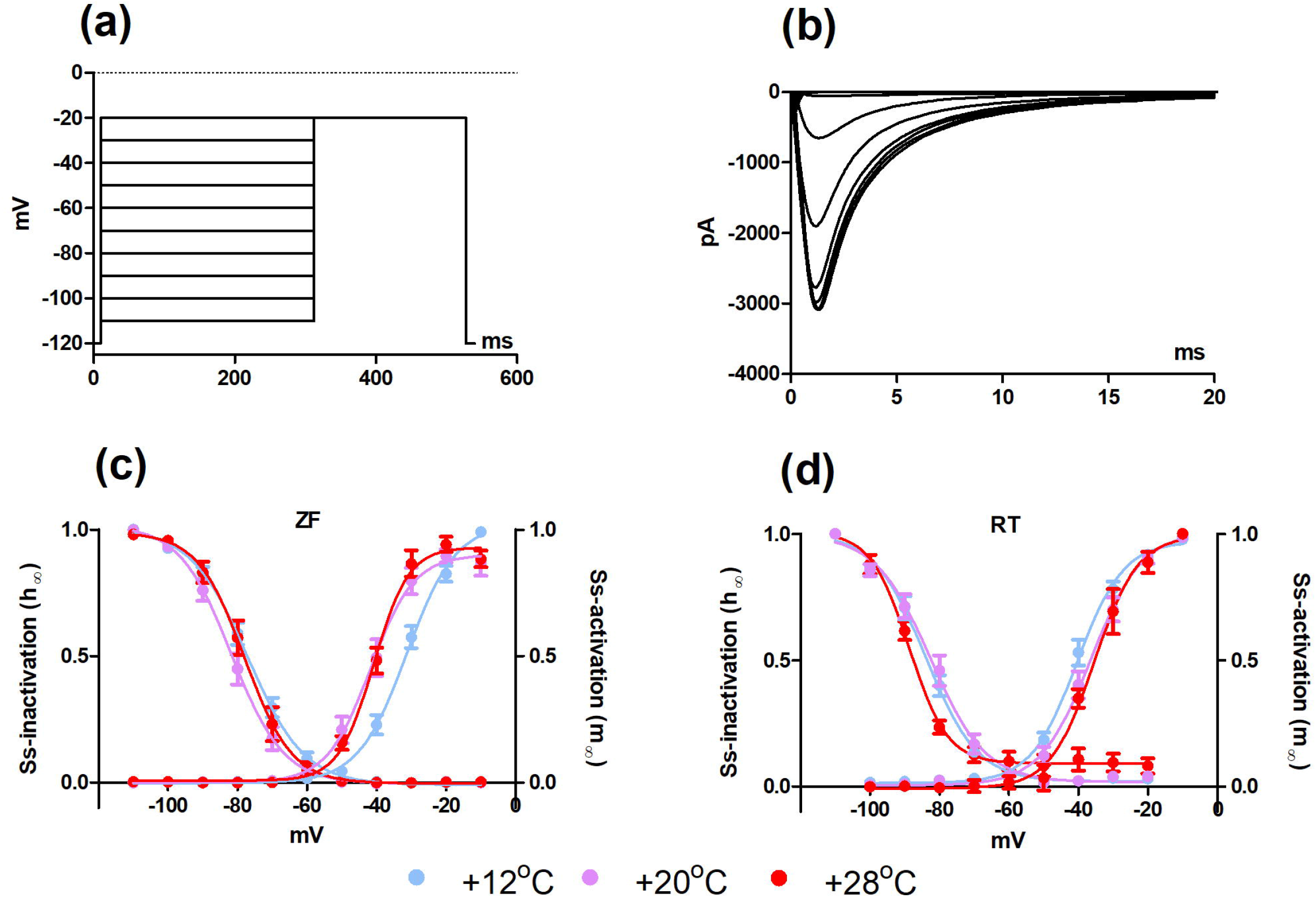
Voltage- and temperature-dependence of steady-state activation and inactivation of I_Na_ in ventricular myocytes of zebrafish and rainbow trout. (a) The two-step voltage protocol used to elicit I_Na_. (b) A representative recording from a zebrafish ventricular myocyte indicating tracings of I_Na_ at −20 mV following 1-s depolarizations from −120 mV to −110 – +30 mV. (c) Steady-state activation and inactivation of zebrafish ventricular I_Na_ (n=87). (d) Steady-state activation and inactivation of rainbow trout ventricular I_Na_ (n=58). Statistically significant differences (p<0.05) are shown as follows: * 12°C vs. 28°C, & 12°C vs. 20°C, # 20°C vs. 28°C.

**Table 2.** Half-voltage (V_0.5_) and slope factor of steady-state (SS) activation and inactivation for I_Na_ of zebrafish ventricular myocytes and Na_V_1.4 and Na_V_1.5 channels expressed in HEK cells.

|  | SS-inactivation |  | SS-activation |  |
| --- | --- | --- | --- | --- |
| | $V_{0.5}$ (mV) | slope | $V_{0.5}$ (mV) | slope |
| **Ventricular myocytes** |  |  |  |  |
| 12°C | -77.2±1.4<sup>a</sup> | -7.3±0.5<sup>a</sup> | -30.3±1.6<sup>a</sup> | 6.7±0.5<sup>a</sup> |
| 20°C | -80.6±1.8<sup>a</sup> | -5.8±0.4<sup>b</sup> | -40.2±2.3<sup>b</sup> | 4.9±0.3<sup>b</sup> |
| 28°C | -77.6±1.9<sup>a</sup> | -5.3±0.6<sup>b</sup> | -39.6±1.8<sup>b</sup> | 5.1±0.7<sup>b</sup> |
| **Nav1.4** |  |  |  |  |
| 12°C | -83.5±2.1<sup>a</sup> | -5.5±0.8<sup>a</sup> | -35.9±3.2<sup>a</sup> | 8.0±0.8<sup>a</sup> |
| 20°C | -85.0±4.0<sup>a</sup> | -6.5±0.6<sup>a</sup> | -42.4±2.5<sup>a</sup> | 5.4±0.4<sup>b</sup> |
| 28°C | -76.8±2.3<sup>a</sup> | -7.3±1.3<sup>a</sup> | -39.0±3.6<sup>a</sup> | 6.6±0.7<sup>a,b</sup> |
| **Nav1.5** |  |  |  |  |
| 12°C | -76.2±2.1<sup>a</sup> | -7.8±1.5<sup>a</sup> | -41.2±4.1<sup>a</sup> | 7.8±1.0<sup>a</sup> |
| 20°C | -78.3±2.4<sup>a</sup> | -8.5±1.7<sup>a</sup> | -33.4±1.7<sup>a</sup> | 7.0±0.8<sup>a</sup> |
| 28°C | -71.1±1.7<sup>a</sup> | -5.5±0.5<sup>a</sup> | -34.1±2.4<sup>a</sup> | 5.5±0.6<sup>a</sup> |

Statistically significant differences ( $p < 0.05$ ) between temperatures are shown by dissimilar letters.

**Table 3.**
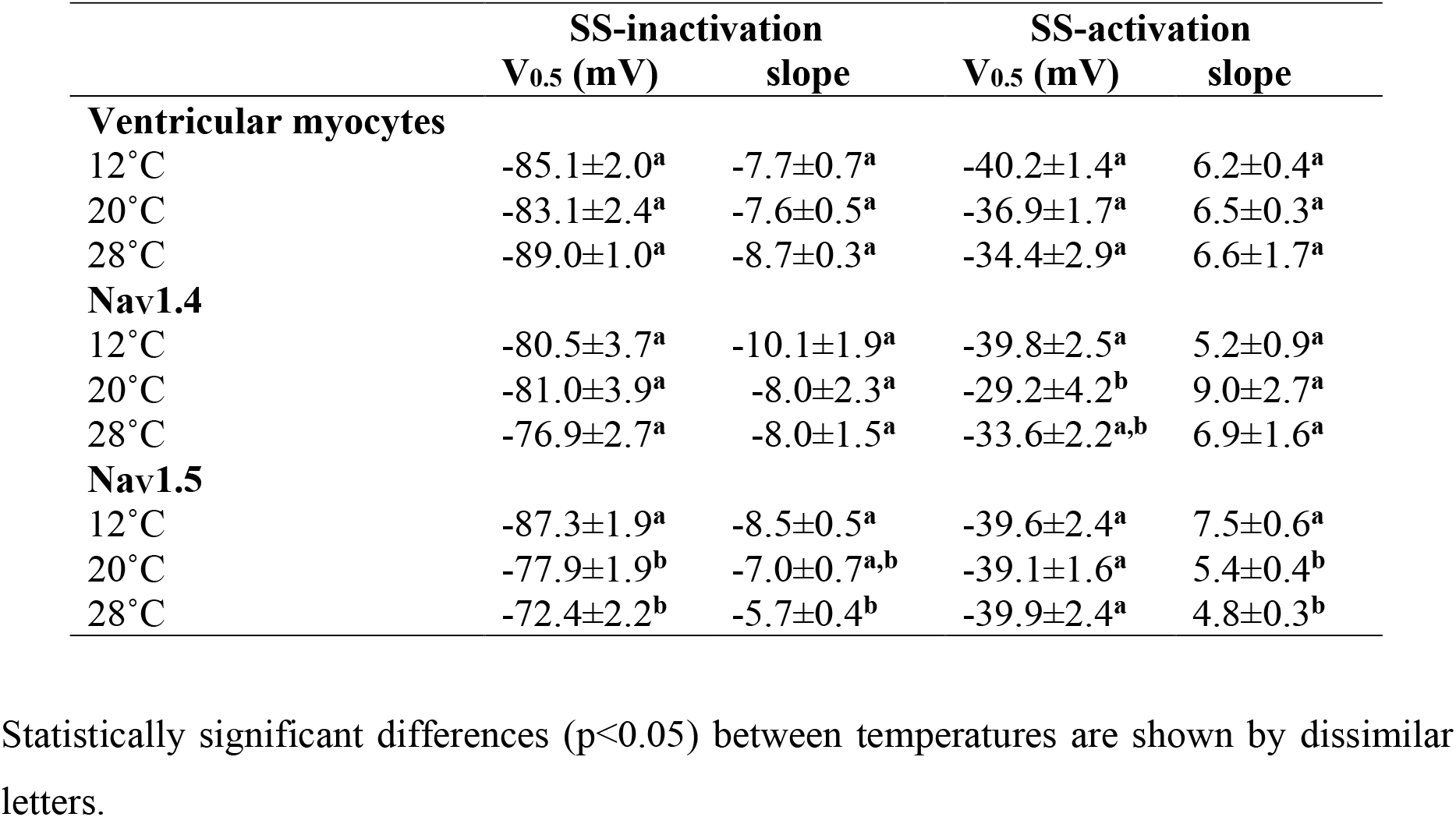
Half-voltage (V_0.5_) and slope factor of steady-state (SS) activation and inactivation for I_Na_ of rainbow trout ventricular myocytes and Na_V_1.4 and Na_V_1.5 channels expressed in HEK cells.

### Temperature-dependence of I_Na_ density and charge transfer generated by Na_V_1.4 or Na_V_1.5 channels in HEK cells

Zebrafish and rainbow trout Na_V_1.4 and Na_V_1.5 Na^+^ channels were expressed in the same cellular environment for a direct comparison between temperature-dependencies of the orthologous gene products (Fig. 4).

**Fig. 4.**
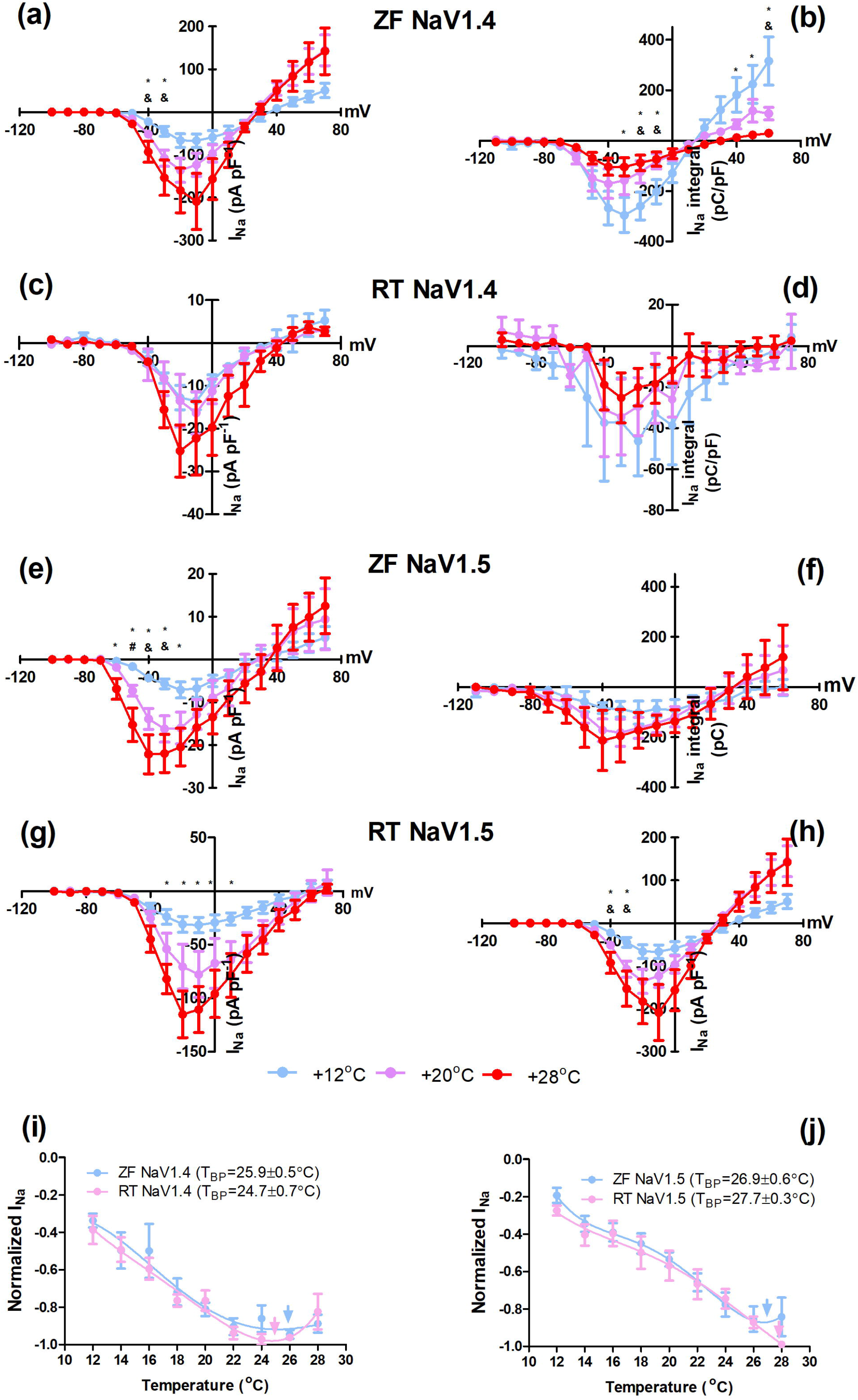
Voltage− and temperature-dependence of current density and charge transfer of I_Na_ generated by zebrafish and rainbow trout Na_V_1.4 and Na_V_1.5 channels in HEK cells. (a-d) Mean results (± SEM) for current-voltage relationship (a, c) and charge transfer (b, d) of I_Na_ at three different temperatures for zebrafish (n=49) (a, b) and rainbow trout (n=15) (c, d) Na_V_1.4 channels. (e-h) Mean results (± SEM) for current-voltage relationship (e, g) and charge transfer (f, h) of I_Na_ at three different temperatures for zebrafish (n=30) (e, f) and rainbow trout (n=27) (g, h) Na_V_1.5 channels. (i and j) Current density (i) and charge transfer (j) of I_Na_ in heat ramp experiments with zebrafish and rainbow trout Na_V_1.4 (i) and Na_V_1.5 (j) channels. Number of tested cells are shown in the figure. Statistically significant differences (p<0.05) are shown as follows: * 12°C vs. 28°C, & 12°C vs. 20°C, # 20°C vs. 28°C.

I_Na_ generated by the zebrafish Na_V_1.4 channels responded to acute temperature increases in similar manner as the native ventricular I_Na_: the current density increased, and the charge transfer decreased with increasing temperature (Fig. 4a, b). I_Na_ generated by the trout Na_V_1.4 channels did not show any statistically significant differences between the 3 temperatures for either current density or charge transfer (Fig. 4c, d) (The expression level of trout Na_V_1.4 channels in HEK cells was much lower than zebrafish Na_V_1.4 channels, which may have reduced the resolution of the analysis). The I_Na_ generated by trout Na_V_1.4 channels was, however, much more heat tolerant than the native trout ventricular I_Na_: current density and charge transfer strongly increased with increasing temperature in HEK cells, while in ventricular myocytes both variables were strongly reduced at 28°C.

I_Na_ density of the zebrafish Na_V_1.5 channels in HEK cells increased with increasing temperature (p<0.05) (Fig. 4e). Differently from the ventricular I_Na_, the heterologously expressed Na_V_1.5 channels positively responded to warming (at −50 mV) up to 28°C i.e. I_Na_ indicated a better heat tolerance in HEK cell membranes than in ventricular myocytes. No statistically significant differences were found in charge transfer by the heterologously expressed Na_V_1.5 channels (Fig. 4f). I_Na_ density of the rainbow trout Na_V_1.5 channels in HEK cells increased with warming from 12°C to 28°C (Fig. 4g). This is a striking deviation from the thermal response of the ventricular I_Na_, which was strongly depressed at 28°C. Charge transfer by rainbow trout Na_V_1.5 channels was significantly increased by acute warming in HEK cells, in contrast to the observations of ventricular I_Na_ (Fig. 4h).

### Inactivation kinetics of I_Na_ generated by Na_V_1.4 and Na_V_1.5 channels in HEK cells

The inactivation rate of I_Na_ increased with increasing temperature and with membrane depolarization. There were marked gene-specific differences in the rate of I_Na_ inactivation (Fig. 5). The rate of inactivation of I_Na_ produced by Na_V_1.5 channels was much slower than that produced by Na_V_1.4 channels for both zebrafish and trout genes (p<0.05). In contrast to gene-specific differences, interspecies differences in inactivation rate of I_Na_ were non-existent or small. The time constant of I_Na_ inactivation of Na_V_1.5 channels was almost identical for zebrafish and trout genes. In contrast, the I_Na_ generated by Na_V_1.4 channels was faster for zebrafish than trout genes at more negative voltages (at −30 mV at 20°C and at −30 and −20 mV at 28°C) (Fig 5.d, f).

**Fig. 5.**
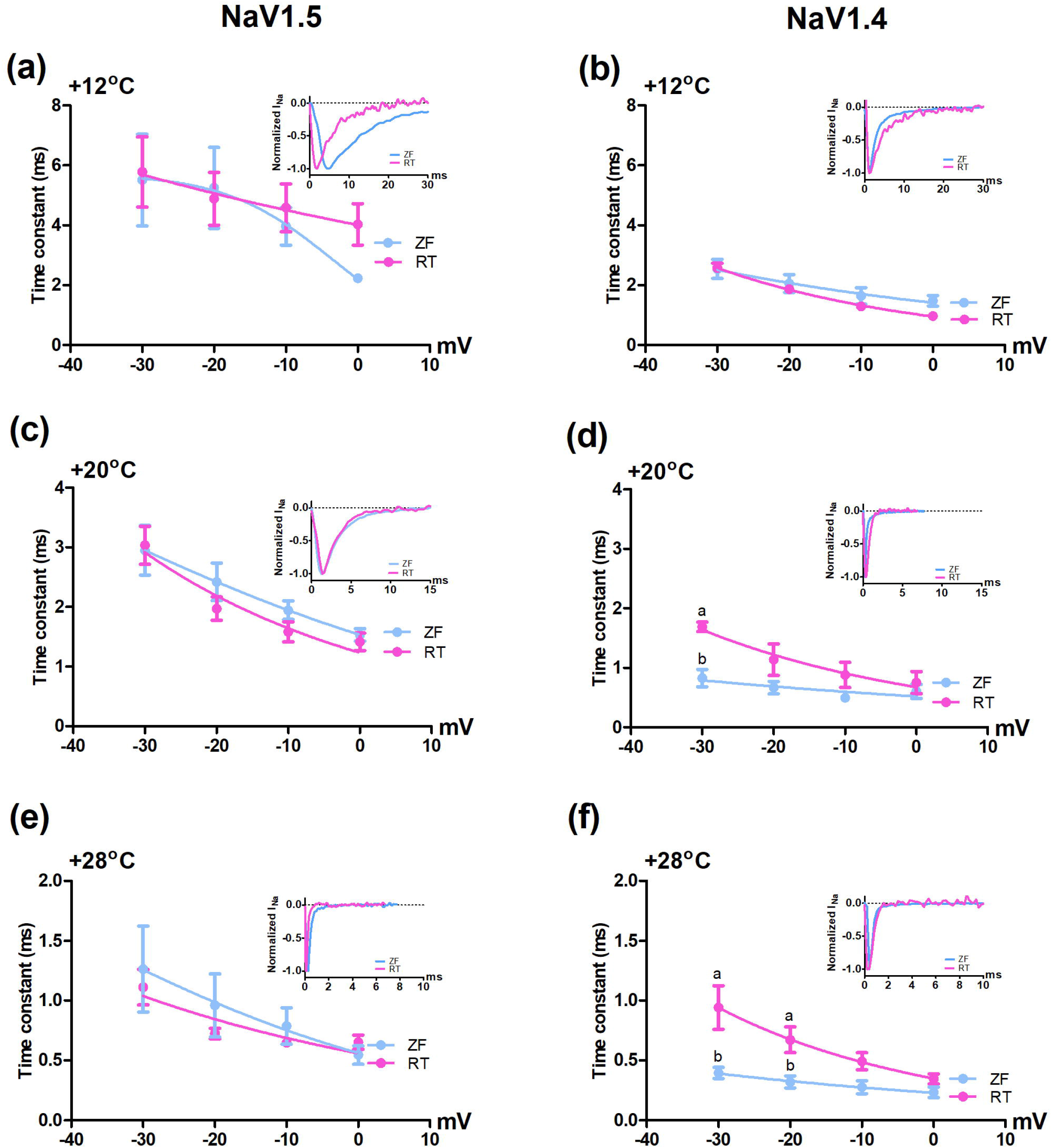
Temperature-dependence of inactivation kinetics of I_Na_ for zebrafish Na_V_1.4 and Na_V_1.5 channels in HEK cells. I_Na_ was elicited from the holding potential of −120 mV to −30 – 10 mV for 30 ms. (a-f) Means (± SEM) of inactivation time constant (τ) for Na_V_1.4 (a, c and e) and Na_V_1.5 (b, d and f) at 12°C, 20°C and 28°C as indicated. The results are from 39 and 42 myocytes from zebrafish and rainbow trout, respectively. Representative tracings of I_Na_ at −30 are shown in the upper right corner of each panel. Statistically significant differences (p<0.05) are shown as follows: * 12°C vs. 28°C, & 12°C vs. 20°C, # 20°C vs. 28°C.

Interestingly, the rate of I_Na_ inactivation of heterologously expressed zebrafish Na_V_1.4 and Na_V_1.5 Na^+^ channels were both much faster than the inactivation kinetics of the endogenous ventricular I_Na_. Opposite to the findings in zebrafish, the I_Na_ inactivation rate of trout Na_V_1.5 channels was much slower than that of trout ventricular myocytes. In contrast, the inactivation rates of I_Na_ for heterologously expressed trout Na_V_1.4 channels and trout ventricular myocytes were similar.

### Steady-state activation and inactivation of I_Na_ generated by Na_V_1.4 and Na_V_1.5 channels in HEK cells

Acute increases in temperature had only weak effects on the voltage-dependence of SS activation and inactivation of I_Na_ generated by the heterologously expressed Na^+^ channels (Tables 1 and 2; Fig. 6). The only change for the zebrafish I_Na_ produced by Na_V_1.5 channels was the reduced slope of SS activation at 20°C and 28°C relative to that at 12°C (p<0.05). Similar changes were observed for the rainbow trout I_Na_ generated by Na_V_1.5 channels (p<0.05) (Fig. 6). In addition, the rainbow trout Na_V_1.5 channels had a positive shift in the voltage-dependence and a decrease in the slope factor of the I_Na_ SS inactivation (p<0.05) (Table 2, Fig. 6h). Elevated temperatures also reduced the slope factor of SS activation of the trout Na_V_1.4 channels (p<0.05) (Table 3, Fig. 6e).

**Fig. 6.**
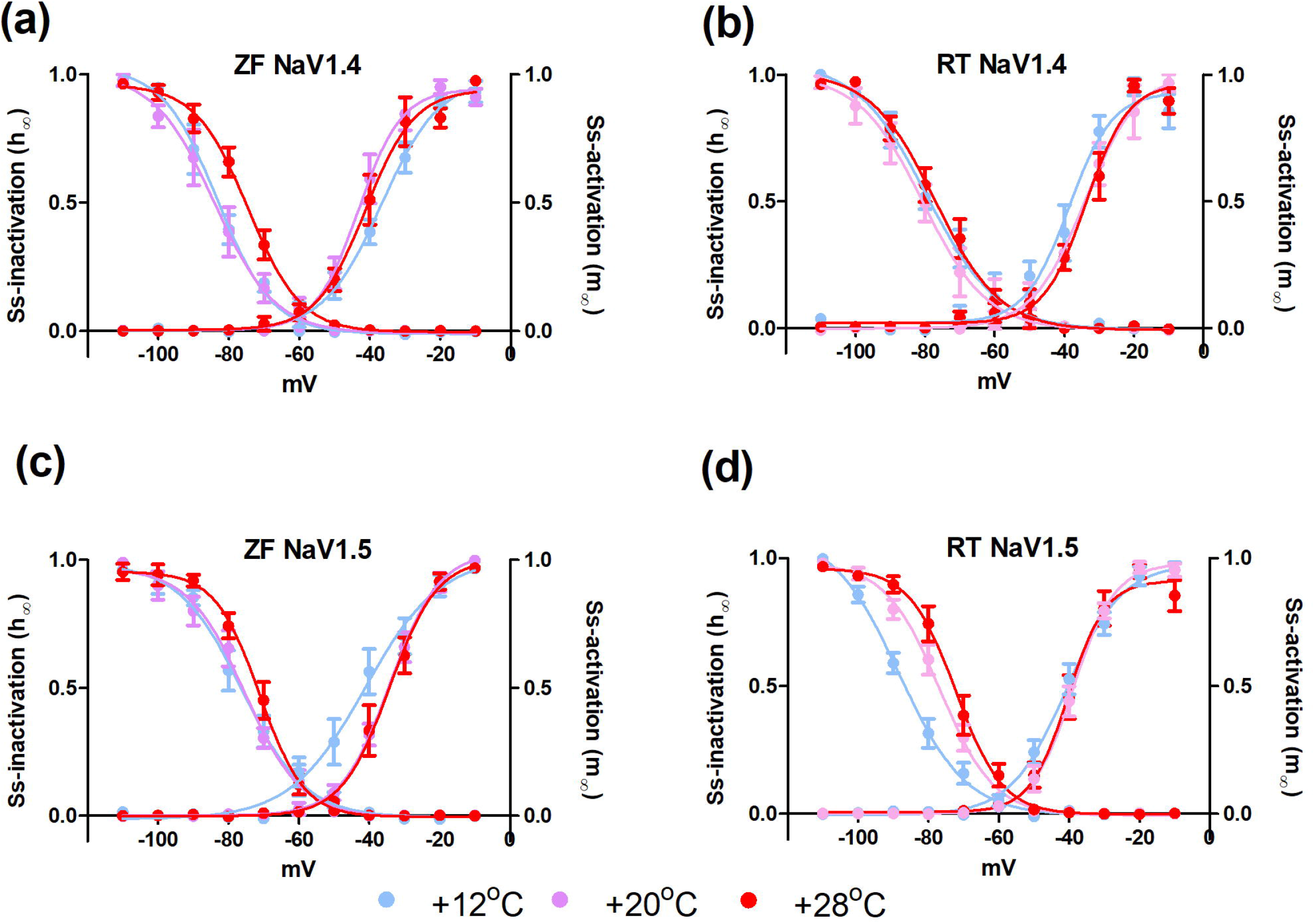
Voltage- and temperature-dependence of steady-state activation and inactivation of I_Na_ for Na_V_1.4 and Na_V_1.5 channels in HEK cells. The voltage protocol was the same as in Fig. 2. (a, b) Steady-state activation and inactivation of zebrafish (n=49) (a) and rainbow trout (n=33) (b) Na_V_1.4 channels. (c, d) Steady-state activation and inactivation of zebrafish (n=49) (c) and rainbow trout (n=50) (d) Na_V_1.5 channels. Statistically significant differences (p<0.05) are shown as follows: * 12°C vs. 28°C, & 12°C vs. 20°C, # 20°C vs. 28°C.

## DISCUSSION

The present results can be summarized in three major findings. (1) The properties of the endogenous I_Na_ of zebrafish and rainbow trout ventricular myocytes differ markedly in terms of heat tolerance and inactivation kinetics with the zebrafish I_Na_ being more heat tolerant and more slowly inactivating. (2) The major Na^+^ channel isoforms of zebrafish and rainbow trout ventricles, Na_V_1.5 and Na_V_1.4 respectively, show only minor interspecies differences, when expressed in HEK cells i.e. the orthologous Na^+^ channel alpha subunits are functionally (heat tolerance, inactivation kinetics) similar in the same membrane matrix. (3) When expressed in HEK cells I_Na_ generated by Na_V_1.4 and Na_V_1.5 isoforms of both species show large channel-specific differences in inactivation kinetics with Na_V_1.4 being fast and Na_V_1.5 slow. Taken together, the species-specific properties of ventricular I_Na_ seem to be determined partly by the expressed Na^+^ channel alpha subunit - Na_V_1.5 in zebrafish and Na_V_1.4 in rainbow trout – and partly by the biophysical properties of the lipid matrix. Notably, the better heat-tolerance of I_Na_ seems to be related to the slower inactivation kinetics of Na^+^ channels.

The endogenous I_Na_ of zebrafish and rainbow trout ventricular myocytes have very different inactivation kinetics: the I_Na_ of zebrafish ventricular myocytes inactivates much more slowly than I_Na_ of rainbow trout ventricular myocytes. At 12°C the inactivation time constant of the zebrafish I_Na_ was almost an order of magnitude bigger than that of the rainbow trout (5.0 ± 0.7 ms vs. 0.5 ± 0.03 ms at −20 mV). This difference largely, but not completely, disappeared when the rate of I_Na_ inactivation was measured at the acclimation temperatures of the fish (0.8 ± 0.09 ms for zebrafish at 28°C vs 0.5 ± 0.03 ms for trout at 12°C). Since thermal acclimation does not have any effect on the inactivation kinetics of the rainbow trout ventricular I_Na_ (Haverinen and Vornanen, 2004), the differences in the inactivation rate between the two species can be regarded as adaptations of I_Na_ to the respective habitat temperatures of the species. In general, electrical excitation in nerves and muscle tissues of ectothermic animals is adapted to work best at lower temperatures in comparison to the tissues of the endothermic animals. For instance, at the typical mammalian body temperatures (36°C-38°C) the AP conduction of ectothermic nerve fibres suffers from heat block (Hodgkin and Katz, 1949; Volgushev et al., 2000). The gating kinetics of the plasma membrane ion channels are probably responsible for the thermal adaptation of AP conduction, allowing APs to propagate at the proper rate and frequency at the typical habitat temperatures of the species. Because zebrafish live at warmer habitats than rainbow trout, the slow inactivation kinetics - when measured at the same experimental temperature - of its ventricular I_Na_ may provide better excitability at higher temperatures than the fast inactivating I_Na_ of the rainbow trout. However, when the intrinsic heart rates of the two species are compared at their acclimation temperatures (130 bpm for zebrafish vs. 60 bpm for rainbow trout) (Aho and Vornanen, 2001; Vornanen and Hassinen, 2016), the rate of I_Na_ inactivation still seems to be slightly slower in zebrafish than rainbow trout.

The slow rate of I_Na_ inactivation in zebrafish ventricular myocytes may protect against heat-induced impairment of electrical excitability (Park et al., 2016; Touska et al., 2018). This is consistent with our hypothesis that Na^+^ channels with slow gating kinetics better maintain electrical excitability at high temperatures (Vornanen, 2020). Indeed, findings from the pain receptors of human skin strongly suggest that the slow gating kinetics of Na^+^ channels are needed for heat resistance of I_Na_ (Touska et al., 2018). At the mammalian body temperature, most Na^+^ channel isoforms operate close to their optimum and a slight increase of temperature above the typical body temperature limits the density and charge transfer of I_Na_ (Touska et al., 2018). However, I_Na_ of the mammalian pain receptors works well at temperatures much above the body temperature (43°C-50°C) largely owing to the thermal properties of the NaV1.9 Na^+^ channel isoform. Inactivation kinetics of this channel are more than an order of magnitude slower than those of NaV1.1-NaV1.8 isoforms (Balbi et al., 2017; Touska et al., 2018). Due to the slow inactivation of NaV1.9 it provides enough charge for depolarization of the plasma membrane at high temperatures. Analogously, the slow inactivation rate of the zebrafish cardiac I_Na_ provides more inward charge for the same peak amplitude of I_Na_ or the same number of active Na^+^ channels than the fast inactivating trout cardiac I_Na_ (Fig. 4). Indeed, the charge transfer/peak current ratio of the zebrafish I_Na_ is much higher at 28°C than that of the rainbow trout I_Na_ at 12°C.

The species-specific difference in the inactivation kinetics of the ventricular I_Na_ is partly explained by the difference in the alpha subunit composition of the Na^+^ channels. In zebrafish ventricular myocytes, the main alpha subunit is Na_V_1.5 which at the transcript level represents 83.1% of all ventricular Na^+^ channels. Transcripts of the Na_V_1.4 consist only 16.2% of the zebrafish ventricular Na^+^ channels (Haverinen et al., 2018). In the ventricle of the rainbow trout, the situation is opposite: Na_V_1.4 channels form 80% of all Na^+^ channel transcripts, while Na_V_1.5 channels represent only 20% of the total channel population (Haverinen et al., 2007). Thus, in zebrafish ventricle Na^+^ channels are mainly of the slow isoform, while in the ventricle of the rainbow trout they are largely composed of the fast isoform. Based on their tissue distribution in mammals, Na_V_1.5 and Na_V_1.4 are often called as ‘cardiac’ and ‘skeletal’ isoform, respectively (Zimmer et al., 2015). The mammalian Na_V_1.4 is kinetically faster than Na_V_1.5 and therefore functionally better to elicit fast twitches at high frequencies in skeletal muscle fibres (Wang et al., 1996; Sheets and Hanck, 1999). Although the classification of Na_V_1.5 and Na_V_1.4 to cardiac and skeletal isoforms, respectively, is not valid for fish, the kinetic similarities between the orthologous mammalian and piscine Na^+^ channels persist. The fast inactivation kinetics of the rainbow trout Na_V_1.4 can be regarded adaptive for heart function of this cold-dwelling fish.

The dominance of Na_V_1.5 alpha subunits in the zebrafish heart only partially explains the slow inactivation kinetics of the ventricular I_Na_. Unlike the rainbow trout, where the inactivation rate of Na_V_1.4 channels in HEK cells relatively closely matches with the inactivation rate of the endogenous ventricular I_Na_, in zebrafish there is a large difference in the inactivation kinetics of the heterologously expressed Na_V_1.5 and the endogenous ventricular I_Na_. Since Na_V_1.4 and Na_V_1.5 Na^+^ channel alpha subunits of zebrafish and rainbow trout share the inactivation kinetics in the heterologous environment of HEK cells, the slow inactivation rate of the zebrafish ventricular I_Na_ is probably partly due to the biophysical properties of the lipid bilayer of ventricular myocytes. Phospholipid bilayer composition of the plasma membrane affects function of the integral membrane proteins mainly in a non-specific manner by its biophysical properties (Lundbaek et al., 2004). Elasticity (stiffness) of the lipid bilayer, determined by the lipid composition, regulates the inactivation of the voltage-gated Na^+^ channels. Decrease in membrane stiffness induced by amphiphiles like Triton-X or β-octyl-glucoside accelerate the rate of I_Na_ inactivation, while increase in membrane stiffness due to elevated cholesterol content decreases the rate of I_Na_ inactivation (Lundbaek et al., 2004). Therefore, it is possible that the lipid composition of the sarcolemma in zebrafish ventricular myocytes differs from that of HEK cells and rainbow trout myocytes, and results in increased stiffness which slows the rate of I_Na_ inactivation.

Although we do not have a complete answer to the slow inactivation rate of the zebrafish ventricular I_Na_, some possible explanations (in addition to the membrane stiffness) can be mentioned to guide future studies. Na^+^ channels are heteromultimers of large pore-forming alpha subunits and small accessory beta subunits. Beta subunits are needed for proper transportation of the alpha subunits into the plasma membrane and they may also affect kinetics of the I_Na_. However, beta subunits are unlikely to cause the slow inactivation of the zebrafish I_Na_ as they tend to enhance the rate of Na^+^ channel inactivation and recovery from inactivation (Chen and Cannon, 1995; Goldin, 2003). A more likely contributing factor is the fibroblast growth factor orthologous factor 2 (FGF2), which binds to the inactivation domain of the C terminus in the Na^+^ channel alpha subunit and strongly slows the rate of inactivation (Liu et al., 2003; Li et al., 2020). Interestingly, the FGF2 knock-out mice are highly sensitive to temperature change and show more cardiac conduction defects when their core body temperature is elevated (Park et al., 2016). Future studies should examine the role of FGFs in temperature-dependence of electrical excitability of the fish heart.

Acute heat challenge experiments indicated that the zebrafish ventricular I_Na_ is much more resistant to high temperatures than the rainbow trout I_Na_. Thus, the heat tolerance of I_Na_ seems to correlate with the upper thermal tolerance of the fish and its heart rate, although in both species the optimum temperature of I_Na_ is lower than the CTmax of the fish and the *T*_BP_ of the intrinsic heart rate (Beitinger, 2000; Aho and Vornanen, 2001; Cortemeglia and Beitinger, 2005; López-Olmeda and Sánchez-Vázquez, 2011; Vornanen and Hassinen, 2016). It is clear that I_Na_ of the trout ventricle would be almost non-functional at the acclimation temperature of the zebrafish, and it is likely that the kinetics of the zebrafish I_Na_ would be too slow for the trout heart at freezing temperatures.

Warming-induced decrease of heart rate is shown to be caused by atrioventricular block, probably due to the reduced excitability of the ventricle (Haverinen and Vornanen, 2020). Therefore, the thermal properties of I_Na_ are likely to affect the species-specific *T*_BP_ of heart rate. Warming-induced decrease of I_Na_ and simultaneous increase in membrane K^+^ leak via I_K1_ results in source-sink mismatch which may prevent AP generation (Vornanen, 2016; Haverinen and Vornanen, 2020; Vornanen, 2020). At the low temperatures there is some excess of I_Na_ relative to the membrane leak via I_K1_ (I_Na_/I_K1_ ≥1.0) called safety factor. When temperature rises the charge transfer of I_Na_ starts to decline (at the species-specific *T*_BP_) due to increased rate of inactivation: the safety factor is lost (I_Na_/I_K1_ <1.0) and excitation fails. In this respect the difference in heat tolerance between the ventricular I_Na_ of the rainbow trout and the I_Na_ generated by the heterologously expressed Na_V_1.4 and Na_V_1.5 channels of the trout may be important. Expression of the trout I_Na_ in the HEK cell membrane makes it much more heat tolerant than it is in the native membrane surroundings. Notably, when zebrafish and trout Na_V_1.4 and Na_V_1.5 were expressed in HEK cells inactivation rate and heat tolerance of I_Na_ were similar for the orthologous channels. It seems that the biophysical properties of the mammalian cell membrane shift the thermal tolerance window of the trout Na^+^ channels to higher temperatures which would, however, be suboptimal at the habitat temperature of the cold-dwelling fish.

### Summary and perspectives

Heat tolerance and inactivation kinetics of I_Na_ differ strongly between zebrafish and rainbow trout ventricular myocytes. In contrast, heat tolerance and inactivation kinetics of Na_V_1.4 and Na_V_1.5 channels of zebrafish and rainbow trout are similar when expressed in the same lipid matrix of HEK cells. Species-specific thermal adaptation of the ventricular I_Na_ is largely achieved by expressing a specific alpha isoform subunit of Na^+^ channel: the slowly inactivating Na_V_1.5 in zebrafish which tolerate well high temperatures and the fast inactivating Na_V_1.4 in rainbow trout which favour cold waters. Differences in elasticity (stiffness) of the lipid bilayer may be also involved in thermal adaptation of I_Na_. These findings suggest that both the protein component and its lipid matrix are involved in thermal adaptation of the voltage-gated Na^+^ channels of the fish heart. Future studies should examine the extent to which these components are flexible under temperature acclimation and therefore able to accommodate the electrical excitability of the heart for seasonal temperature changes and peak summer temperatures. Since electrical excitability is regulated in basically the same way in all excitable cells, studying the biophysical properties of neuronal and muscular I_Na_ could provide cues to its role in thermal homeostasis/death of ectotherms.

## List of symbols and abbreviations

AP: action potential
HEK cell: human embryonic kidney cell
I_K1_: inward rectifier potassium current
I_Na_: sodium ion current
Na_V_1.4: sodium channel alpha subunit 1.4
NaI_V_1.5: sodium channel alpha subunit 1.5
Q_10_: thermal coefficient
SCN: sodium channel
*T*_BP_: breakpoint temperature
V_rest_: resting membrane potential
V_th_: threshold potential

## Acknowledgements

We would like to thank Kontiolahti fish farm for providing the trout. Anita Kervinen is acknowledged for taking care of the fish at the university and preparing solutions for experiments.

## Author contributions

Conceptualization: M.V.; Methodology: J.H., I.D., M.H.; Investigation: J.H., I.D., D.V.A., M.H.; Writing -original draft: J.H., D.V.A., M.H.; Writing -review & editing: M.V.; Visualization: J.H., I.D.; Project administration: M.V.; Funding acquisition: M.V.

